# The lipid transfer proteins Nir2 and Nir3 sustain phosphoinositide signaling and actin dynamics during phagocytosis

**DOI:** 10.1101/2022.12.21.521447

**Authors:** Mayis Kaba, Amado Carreras-Sureda, Paula Nunes-Hasler, Nicolas Demaurex

**Affiliations:** Department of Cell Physiology and Metabolism University of Geneva, Geneva, 1211, Switzerland; Department of Pathology and Immunology, University of Geneva, Geneva, 1211 Switzerland

**Keywords:** Immunity, Cytoskeleton, Lipid transfer, Membrane contact sites, Cell signaling

## Abstract

Changes in membrane phosphoinositides and local Ca^2+^ elevations at sites of particle capture coordinate the dynamic remodeling of the actin cytoskeleton during phagocytosis. Here, we show that the phosphatidylinositol (PI) transfer proteins PITPNM1 (Nir2) and PITPNM2 (Nir3) maintain PI(4,5)P2 homeostasis at phagocytic cups, thereby promoting actin contractility and the sealing of phagosomes. Nir3 and to a lesser extent Nir2 accumulated in ER cisternae juxtaposed to phagocytic cups when expressed in phagocytic mouse fibroblasts. CRISPR-Cas9 editing of Nir2 and Nir3 genes decreased plasma membrane PI(4,5)P2 levels, store-operated Ca^2+^ entry (SOCE), and receptor-mediated phagocytosis, stalling particle capture at cup stage. Re-expression of either Nir2 or Nir3 restored phagocytosis, but not SOCE, proportionally to the PM PI(4,5)P2 levels. Phagosomes forming in Nir2/3-edited cells had decreased overall PI(4,5)P2 levels but normal periphagosomal Ca^2+^ signals. Nir2/3 editing reduced the density of contractile actin rings at sites of particle capture, causing repetitive low-intensity contractile events indicative of abortive phagosome closure. We conclude that Nir-mediated lipid transfer maintains phosphoinositide homeostasis at phagocytic cups, thereby sustaining the signals that initiate the remodeling of the actin cytoskeleton during phagocytosis.

**Summary statement:** Changes in membrane phosphoinositides coordinate actin remodeling during phagocytosis, but whether lipid transport proteins contribute to this process is not known. Here, we show that the phosphatidylinositol transfer proteins Nir2 and Nir3 are recruited to phagocytic cups and drive the formation of contractile actin rings during particle engulfment. Using gene editing and re-expression, we show that Nir2 and Nir3 maintain PI(4,5)P2 signaling competence at phagocytic cups and promote the actin-dependent sealing of phagocytic vacuoles. These observations establish that lipid transport proteins maintain the phosphoinositide signals that drive the remodeling of the actin cytoskeleton during phagocytosis.

## Introduction

Phagocytosis, the engulfment of large particles by cells, is an evolutionarily conserved cellular process required to eliminate invading pathogens and to maintain tissue homeostasis (Flannagan et al., 2012). Particle recognition is mediated by surface receptors for immunoglobulins or complement fragments coating invading pathogens and by receptors for pathogen-associated sugars (Uribe-Querol and Rosales, 2020). The engagement of phagocytic receptors initiates a dramatic remodeling of the plasma membrane accompanied by acute changes in phosphoinositide composition at sites of particle capture. Sequential fluctuations in the local concentration of phosphatidylinositol mono, bis, and tris phosphate regulate distinct trafficking and signaling events during the formation of phagocytic vacuoles and their subsequent fusion with endolysosomes (Botelho et al., 2000; Levin-Konigsberg et al., 2019; Montaño-Rendón et al., 2022), reviewed in (Bohdanowicz and Grinstein, 2013; Flannagan et al., 2012). A spatially restricted transient PI(4,5)P2 elevation occurs at phagocytic cups coinciding with the transient recruitment of the 5-kinase that converts PI(4)P into PI(4,5)P2 (Botelho et al., 2000). The PI(4,5)P2 elevation promotes actin polymerization at phagocytic cups by activating actin nucleators while inhibiting actin severing proteins (Bohdanowicz and Grinstein, 2013; Yeung et al., 2006) As the phagosome seals, PI(4,5)P2 is phosphorylated by the phosphatidylinositol 3-kinase (PI3K) into PI(3,4,5)P3, which in turn becomes transiently enriched at phagocytic cups (Marshall et al., 2001) before being converted to PI(3,4)P2 by phosphoinositide 5-phosphatases (Montaño-Rendón et al., 2022).

Following receptor stimulation, PI(4,5)P2 is also converted by phospholipase C gamma (PLCγ) into diacylglycerol (DAG) and inositol trisphosphate (InsP3). DAG is further converted to phosphatidic acid (PA) by phosphorylation at the PM (Balla, 2013; Cockcroft and Garner, 2011) while InsP3 promotes the release of Ca^2+^ from endoplasmic reticulum (ER) stores. Store depletion activates the Ca^2+^-sensing protein STIM1 that traps and gate the ORAI family of Ca^2+^ channels at ER-PM and ER-phagosomes membrane contact sites (MCS) further stabilized by junctate (Guido et al., 2015; Nunes et al., 2012; Westman et al., 2019). The combined activity of PLCγ and PI3K depletes PI(4,5)P2, promoting actin disassembly at the base of phagocytic cups (Scott et al., 2005) while periphagosomal Ca^2+^ elevations drive the activity of Ca^2+^-dependent actin-severing proteins around phagocytic vacuoles (Nunes et al., 2012). This signaling cascade is required for successful target internalization, and decreasing PI(4,5)P2 levels or preventing periphagosomal Ca^2+^ elevations impair phagocytosis (Botelho et al., 2000; Coppolino et al., 2002; Scott et al., 2005; Nunes et al., 2012). The signals initiated at cups implies that sufficient levels of lipid precursors are delivered at sites of forming phagosomes to fuel the sequential changes in signaling lipids, but the mechanism(s) ensuring the supply of phosphoinositides at sites of particle capture are unknown.

Nir2 (PITPNM1) and Nir3 (PITPNM2) are mammalian homologues of *Drosophila* retinal degeneration protein B (rdgB) harboring an N-terminal PI transfer domain (PITP) that drives the non-vesicular exchange of ER-bound PI for phosphatidic acid (PA) between the ER and target membranes (reviewed in (Balla, 2018). Nir2 and Nir3 are recruited to ER-PM contact sites upon receptor stimulation via interactions with the FFAT motif of the ER-resident vesicle-associated membrane-associated proteins VAP-A and VAP-B (Amarilio et al., 2004; Selitrennik and Lev, 2016). Nir2 maintains PI(4,5)P2 levels at the PM during signaling of Gq-coupled receptors by exchanging PI from the ER for PA on the PM (Chang & Liou, 2015; Kim et al., 2015), thereby preserving Ca^2+^ signaling competence when phosphoinositides are rapidly consumed by PLC. This homeostatic function is clinically relevant as increased Nir2 expression correlates with enhanced epithelial-mesenchymal transition and poor patient prognosis (Keinan et al., 2014). During store-operated Ca^2+^ entry (SOCE) Nir2 is dynamically recruited to STIM/ORAI contact sites by the concomitant Ca^2+^-dependent recruitment of extended synaptotagmin-1 (E-Syt1), which stabilizes ring-shaped ER-PM contact sites at a reduced gap distance (Chang and Liou, 2016; Kang et al., 2019). Whether Nir proteins mediate lipid transfer at phagosomes is not known, but another lipid transfer protein, ORP1L, interacts with VAP proteins at ER-phagosome contact sites and contributes to phagolysosome resolution by transferring the PI(4)P accumulating in late phagosomes to the ER (Levin-Konigsberg et al., 2019).

Given the importance of phosphoinositide signals as a crucial regulator of phagocytosis, we investigated the contribution of Nir-mediated non-vesicular lipid transfer in phagosome formation and maturation. Using correlative light-electron microscopy we show that Nir3 is recruited to membrane contact sites at phagocytic cups. Using GFP fused to plekstrin-homology (PH) domains of PLC (Várnai et al., 1999) or LifeAct-mCherry, we quantify PI(4,5)P2 levels and actin dynamics during FcR-mediated uptake of solid particles by cells lacking or re-expressing Nir2/3 proteins. Depletion of Nir2/3 decreased periphagosomal PI(4,5)P2 and F-actin accumulation around forming phagosomes, stalling phagocytosis at cup stage, without perturbing the subsequent PI(4,5)P2 and Ca^2+^ elevations around successfully internalized particles. This suggests that Nir-mediated lipid transfer occurs at cups and maintains PI(4,5)P2 levels required for the formation of contractile actin rings during phagosome.

## Results

### Nir2 and Nir3 localize to phagocytic cups

The two proteins Nir2 and Nir3 transfer PI from the ER to the PM to replenish PI(4,5)P2 levels following receptor-induced hydrolysis (Chang and Liou, 2015), but whether lipid transport by these proteins contribute to the phagocytic process is unknown. To assess if Nir proteins are recruited to phagocytic vacuoles, we expressed EGPF-tagged Nir2 or Nir3 in COS7 cells rendered phagocytic by expression of the immunoglobulin receptor FcγRIIA-c-myc (Guido et al., 2015) and assessed the location of the fluorescent proteins by confocal imaging. EGFP-Nir2 and to a larger extent EGFP-Nir3 accumulated at sites of particle capture, forming ring structures surrounding internalized particles, indicating that the lipid transfer proteins are recruited to phagocytic cups (Fig. 1A). Since Nir proteins interact with ER-bound FFAT-binding proteins, we tested wither the latter are present at cups by co-expressing EGFP-Nir3 with mCherry-VAP-B, which colocalizes with OPR1L on late phagosomes (Levin-Konigsberg et al., 2019). EGFP-Nir3 colocalized extensively with co-expresed mCherry-VAP-B at cups (Fig. 1B), suggesting that the two proteins are co-recruited to membrane contact sites at cup stage. To establish that Nir3 populates ER-phagosome contact sites, we then performed correlative light-electron microscopy (CLEM). Phagocytic COS7 cells were transfected with EGFP-Nir3 and allowed to ingest opsonized beads for 30 min. Cells were then fixed and imaged by confocal microscopy to locate Nir3-decorated phagosomes (Fig. 1C, left panel). EM tomograms of the labeled phagosomes were then acquired at nanometric resolution. Alignment of the confocal and EM images revealed that EGFP-Nir3 signals coincided with ER membranes around phagocytic cups (Fig 1C, right). These data establish that Nir3 is recruited together with ER-bound proteins to membrane contact sites at phagocytic cups.

**Figure 1.**
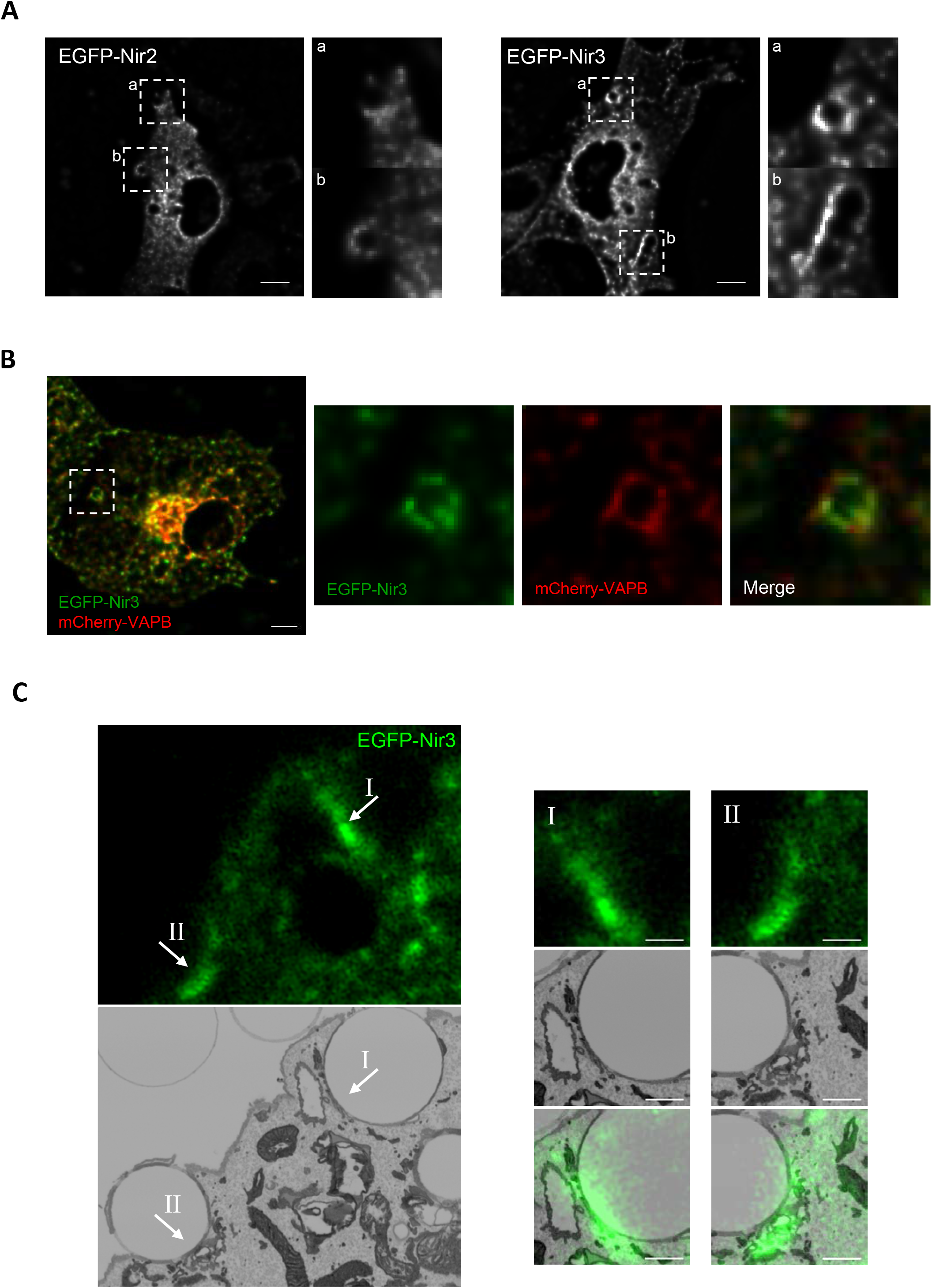
Nir2 and Nir3 are recruited to ER-phagosome contact sites. A) Deconvolved confocal sections of Cos7 cells expressing myc-FcRIIA together with EGFP-Nir2 (left) or EGFP-Nir3 (right) after 30 min of phagocytosis. B) EGFP-Nir3 (green) was co-expressed with mCherry-VAP-B (red). Insets show the phagosome-associated fluorescence. Scale bars: 10 μm. C) Correlative light-electron microscopy (CLEM) images of Cos7 cells expressing myc-FcRIIA and EGFP-Nir3 (green) after 30 min of phagocytosis. Numbered arrows indicate EGFP-Nir3 enriched ER-phagosome contact sites. Scale bars: 1 μm.

### Nir2/3 depletion decreases plasma membrane PI(4,5)P2 levels and SOCE

To assess the role of PI transfer proteins during phagocytosis, we used CRISPR-Cas9 editing to disrupt the *Pitpnm1* and *Pitpnm2* genes coding for Nir2 and Nir3 in mouse embryonic fibroblasts (MEF). Proper editing was validated by the failure of primers targeting the modified genomic regions to amplify a PCR product (Fig. S1A). Real time qPCR with primers targeting non-edited regions indicated that Nir2 and Nir3 mRNA levels were decreased by 80% and 60 %, respectively (Fig. S1B) with similar melting points, indicating a decreased stability of the modified mRNAs. We then measured basal PM PI(4,5)P2 levels with PH-PLCδ1-GFP (Balla and Várnai, 2002) in control and edited cells to verify the functional impact of these genetic manipulations. An intense PH-PLCδ1-GFP signal was detected at the edge of control cells while a weaker and discontinuous signal delineated cells bearing each of the singly or doubly edited gene (Fig. 2A). Quantification of the GFP signal at the cell edge and cytosol revealed that the PM-to-cytosol intensity ratio was significantly reduced in Nir2- and Nir2/3-edited cell lines (Fig. 2B), indicating that depletion of Nir isoforms decreases basal PI(4,5)P2 levels. We then tested whether SOCE, which is mediated by the recruitment of STIM proteins to PI(4,5)P2-rich domains maintained by Nir2 at ER-PM contact sites (Chang & Liou, 2015), was impacted by Nir2/3 knock-down. As expected, the amplitude of the Ca^2+^elevations evoked by the readmission of Ca^2+^ to cells treated with the SERCA inhibitor thapsigargin was severely impacted in Nir2/3-edited cells (Fig. 2C). Stable expression of RFP-tagged Nir2 or Nir3 restored basal PM PI(4,5)P2 levels in Nir2/3-edited cells (Fig. 2D-E and S2A), suggesting that the two lipid transport proteins complement each other for the maintenance of basal PM phosphoinositide levels. Interestingly, SOCE was not restored to WT levels by exogenous overexpression of the tagged Nir proteins (Fig. S2B). These data indicate that depletion of the PI transfer proteins Nir2 and Nir3 decreases basal PM PI(4,5)P2 levels and SOCE in mouse fibroblasts and validate Nir2/3-edited cells as a useful tool to study the role of these transfer proteins.

**Figure 2.**
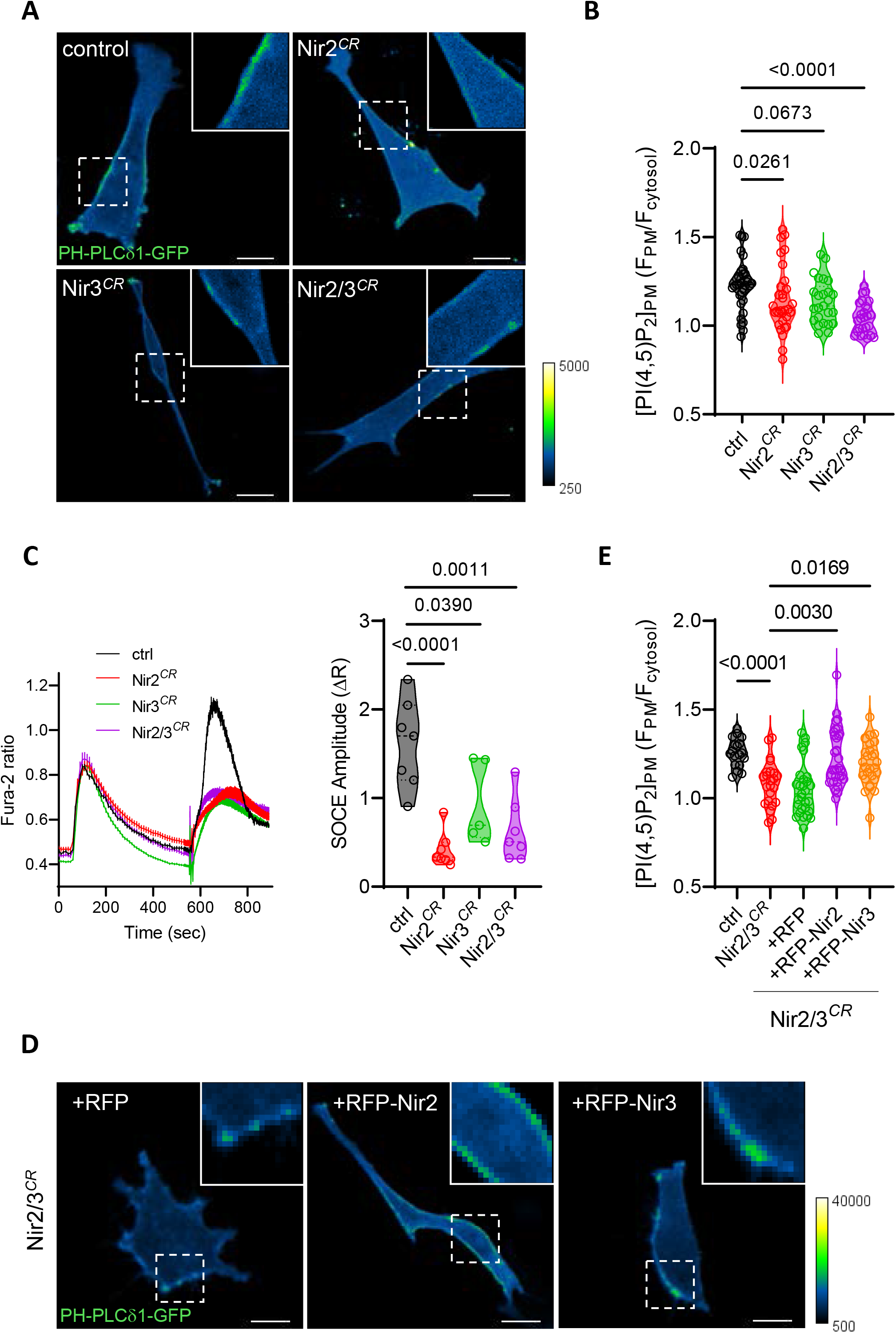
Nir2 and Nir3 sustain PI(4,5)P2 and Ca^2+^ signaling at the plasma membrane. A) Confocal sections of cells expressing PH-PLCδ1-GFP. Insets show PM-associated fluorescence. Scale bars: 10 μm. B) quantification of PM vs. cytosolic fluorescence intensity in control (n=34), Nir2-edited (n=31), Nir3-edited (n=25) and Nir2/3-edited cells (n=26). Kruskal-Wallis test for multiple comparison of individual values across n=3-5 independent experiments. C) left: averaged Ca^2+^ responses evoked by the addition of thapsigargin and 1 mM Ca^2+^ to CRISPR control and Nir2/3-edited cells. Right: Peak amplitude of the response evoked by Ca^2+^readmission. Two-tailed unpaired t test vs. control of N=5-7 independent recordings totaling 152-219 cells. D) Confocal sections of Nir2/3-edited cells expressing PH-PLCδ1-GFP and the indicated constructs. Scale bars: 10 μm. E) quantification of PM vs. cytosolic fluorescence intensity in control (n=18), Nir2/3-edited cells (n=23) and Nir2/3-edited cells expressing RFP (n=29), RFP-Nir2 (n=32), and RFP-Nir3 (n=28). Kruskal-Wallis test for multiple comparison of individual values across n=3-5 independent experiments.

### Nir2/3 depletion decreases phagocytosis

Phosphoinositides are central regulators of the trafficking and signaling events driving phagocytosis. To test whether the depletion of Nir proteins impacts the phagocytic process, we quantified the uptake of opsonized polystyrene beads by MEF cells rendered phagocytic by transient FcγRIIA-c-myc expression (Guido et al., 2015). Phagocytic uptake was quantified as the percentage of cells exhibiting one or more associated fluorescent particles on flow cytometry scatter plots (Fig. 3A). Particle uptake was significantly reduced in Nir2/3-edited cells incubated for 30 and 60 min with opsonized particles (Fig. 3A-B), as was the phagocytic index measured by fluorescence imaging (Fig. 3C). Stable re-expression of RFP-tagged Nir2 or Nir3 partially restored phagocytic uptake in Nir2/3-edited cells (Fig. 3D). Of note, the PM PI(4,5)P2 levels measured with PH-PLCδ1-GFP in the different cell lines correlated well with the proportion of phagocytosing cells (Fig 3E). These data indicate that Nir2/3 depletion reduces the efficiency of phagocytosis and that this defect is proportional to the reduction in PI(4,5)P2 levels caused by the loss of these lipid transport proteins.

**Figure 3.**
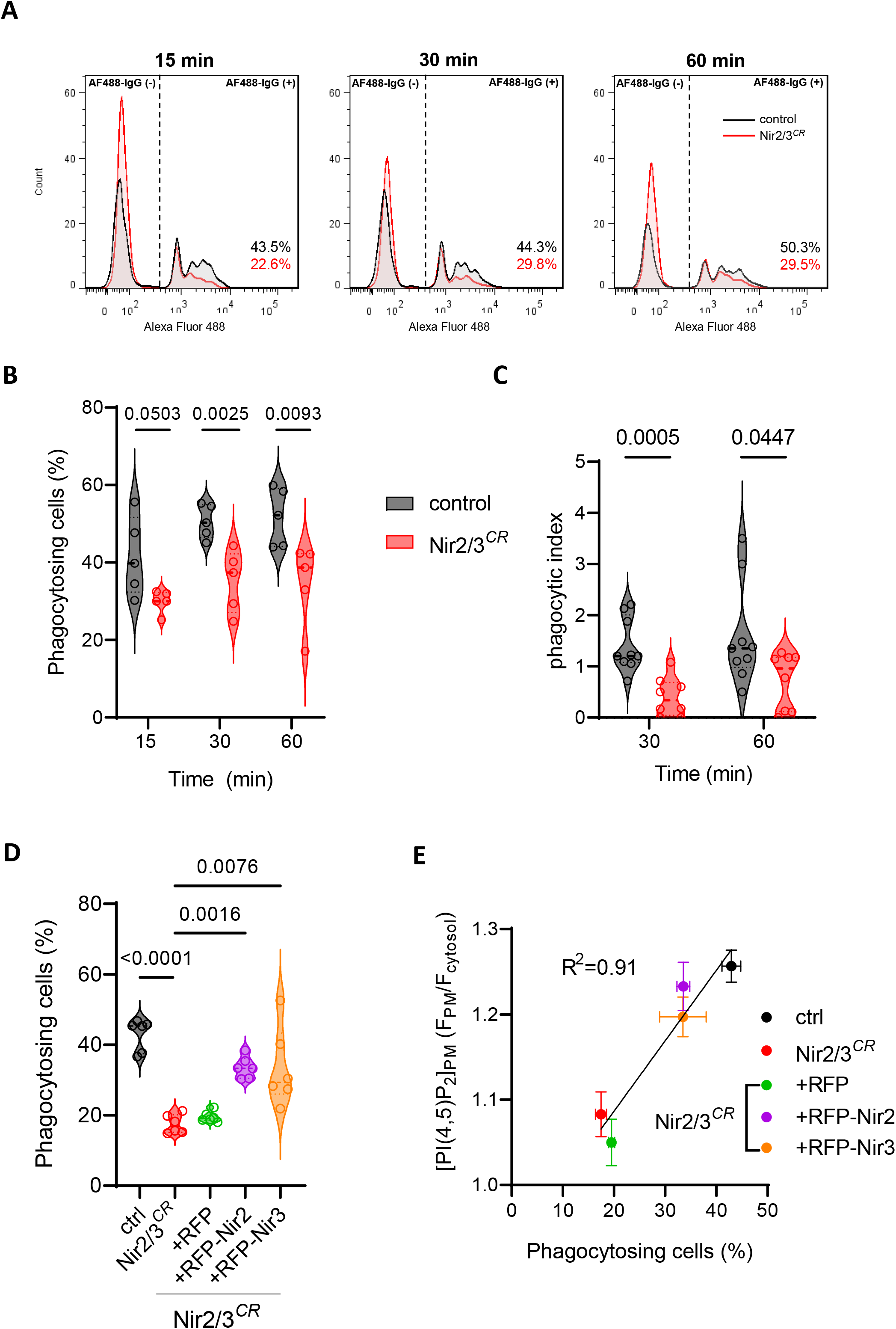
Depletion of Nir2 and Nir3 decreases phagocytosis. A) Flow cytometry fluorescence intensity profiles of control and Nir2/3-edited cells expressing Myc-FcRIIA exposed to IgG opsonized fluorescent particles for the indicated time periods. B) Proportion of cells with phagocytosed particles at the indicated time points. Mean±SEM, n=5, Two-tailed paired t test. C) Proportion of internalized particles in control (n=9) and Nir2/3-edited cells (n=8) measured by microscopy 30 and 60 min after particle exposure. Two-tailed unpaired t test. D) Proportion of cells with phagocytosed fluorescent particles at 60 min. n=6, Two-tailed paired t test for control vs. Nir2/3^*CR*^, Kruskal-Wallis test for multiple comparison for Nir2/3^*CR*^ vs. RFP constructs. E) Correlation between PM PI(4,5)P2 levels measured with PH-PLCδ1-GFP in Fig. 2 and the proportion of phagocytosing cells at 60 min in Panel D. Data are mean±SEM.

### Nir2/3 depletion reduces phagosomal PI(4,5)P2 but not periphagosomal Ca^2+^ signals

Biphasic changes in PI(4,5)P2 levels occur during phagocytosis, with a rapid elevation at sites of particle engagement followed by a decrease as phagosomes seal (Botelho et al., 2000). To establish the role of Nir proteins in phagosomal PI(4,5)P2 dynamics, we imaged local PI(4,5)P2 levels around forming and internalized phagosomes using PH-PLCδ1-GFP. A transient PI(4,5)P2 enrichment was observed around nascent phagosomes 30-90 sec after particle capture (Fig. 4A), both in control and Nir2/3 edited cells. However, quantification of the phagosomal PI(4,5)P2 revealed that while the transient elevation persisted in Nir2/3 edited cells, the local levels of the phosphoinositide were dampened throughout the phagocytic process (Fig.4B). These data indicate that Nir2/3 proteins help sustain high levels of PI(4,5)P2 on the membrane of phagosomes.

**Figure 4.**
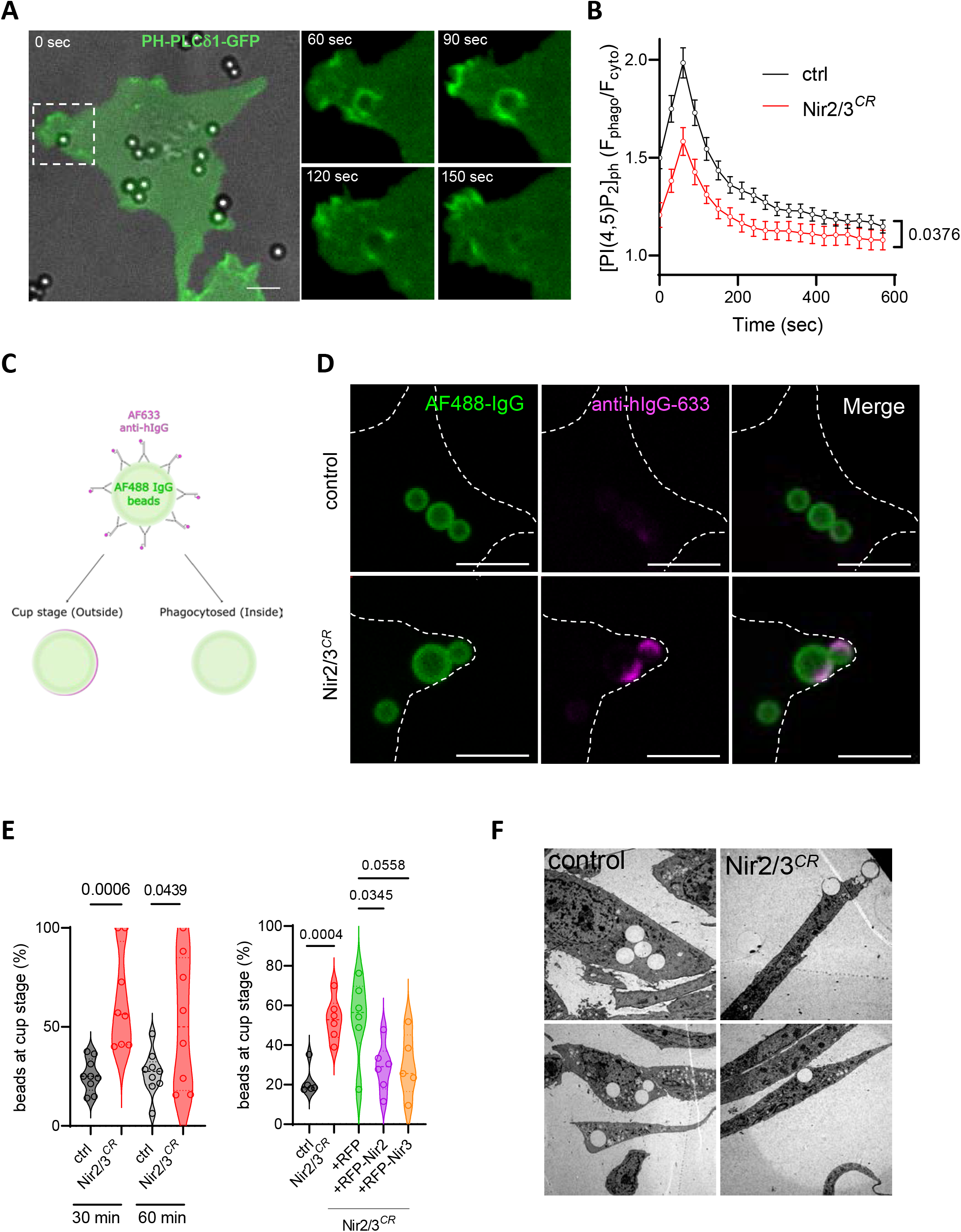
Nir2 and Nir3 depletion reduces phagosomal PI(4,5)P2 and stalls phagocytosis at cup stage. A) Time-lapse images of PH-PLCδ1-GFP dynamics in a control cell phagocytosing IgG-opsonized particles. B) changes in phagosomal vs. cytosolic PH-PLCδ1-GFP fluorescence during particle uptake by control (n=74) and Nir2-edited cells (n=30). Two-way ANOVA with Geisser-Greenhouse correction. Scale bar: 10 μm. C) procedure used to identify non-internalized particles by their accessibility to soluble anti-IgG. Right: Confocal sections of control and Nir2/3-edited cells 30 min after particle exposure. All opsonized particles are coated with fluorescent IgG (green), particles at cup stage are further decorated by anti-IgG (red). Scale bars: 10 μm. E) Left: proportion of particles at cup stage in control (n=9) and Nir2/3-edited cells (n=8) after 30 and 60 min of phagocytosis. Two-tailed unpaired t test of individual values across n=3 independent experiments. Right: Proportion of particles at cup stage in cells expressing or not the indicated constructs. n=6, Two-tailed unpaired t test for ctrl vs. Nir2/3^*CR*^, Kruskal-Wallis test for multiple comparison for RFP vs. RFP-Nir constructs. F) Electron micrographs of control and Nir2/3-edited cells 30 min after particle exposure. Bar 5 μm.

STIM1 is recruited to phagosomes, generating local Ca^2+^ elevations that enhance the efficiency of phacocytosis (Nunes et al., 2012; Nunes-Hasler et al., 2020; Westman et al., 2019). STIM1 binds PI(4,5)P2 via its C-terminal tail and colocalizes with Nir2 at ER-PM contact sites (Kim et al., 2015), prompting us to measure local Ca^2+^ elevations around phagosomes by confocal imaging using Fluo-8 and a low concentration of the Ca^2+^ chelator BAPTA-AM to reduce lateral Ca^2+^ diffusion. Unexpectedly, Ca^2+^ hotspots were detected around 34-40% of (successfully ingested) phagosomes independently of Nir2/3 depletion (Fig. S3), indicating that Nir proteins are dispensable for periphagosomal Ca^2+^ signals.

### Nir2/3 depletion stalls the phagocytic process at the cup stage

We next assessed whether early steps of particle capture were impacted by the loss of Nir2/3 proteins. Cells were allowed to phagocytose opsonized fluorescent beads and the amount of total and surface-bound particles assessed using the bead-associated (green) and solvent-accessible anti-IgG (red) fluorescence signals, respectively (Fig. 4C-D and S4A). The proportion of exposed particles, reflecting beads stuck at cup stage, was significantly increased in Nir2/3-edited cells (Fig. 4E, left). Re-expression of RFP-Nir2 or RFP-Nir3 reduced the proportion of particles stuck at cup stage, linking the defect to reduced Nir-mediated lipid transfer (Fig. 4E, right). Electron microscopy confirmed that most particles were intracellular in control cells and present at cups in Nir2/3-edited cells (Fig. 4F). These results indicate that Nir-mediated lipid transfer facilitates the closure of phagocytic cups.

### Nir2/3 depletion impairs the formation of contractile actin rings during phagocytosis

Since phosphoinositide composition controls actin dynamics during pseudopod formation and phagosome sealing, we next quantified the changes in the density of the cortical actin cytoskeleton forming around captured particles with LifeAct-mCherry. As previously reported, a transient actin enrichment was observed at sites of particle capture immediately after contact, the F-actin probe accumulating into ring structures that rapidly subsided as the particles were internalized (Fig. 5A). Remarkably, multiple rings of actin were observed arising repeatedly for up to 15 min at sites of particle capture in Nir2/3-edited cells (Fig. 5A-B and supplementary video 1). Repetitive ring formation was observed in 32% vs. 4% of the phagocytic events analyzed in Nir2/3-edited and control cells, respectively, a highly significant difference (Fig. 5B, bottom). Moreover, the thickness of the circumferential LifeAct-mCherry signal was reduced in Nir2/3-edited cells (Fig. 5C-D). These data indicate that Nir-mediated lipid transfer facilitates the formation of contractile actin rings during phagocytic uptake.

**Figure 5.**
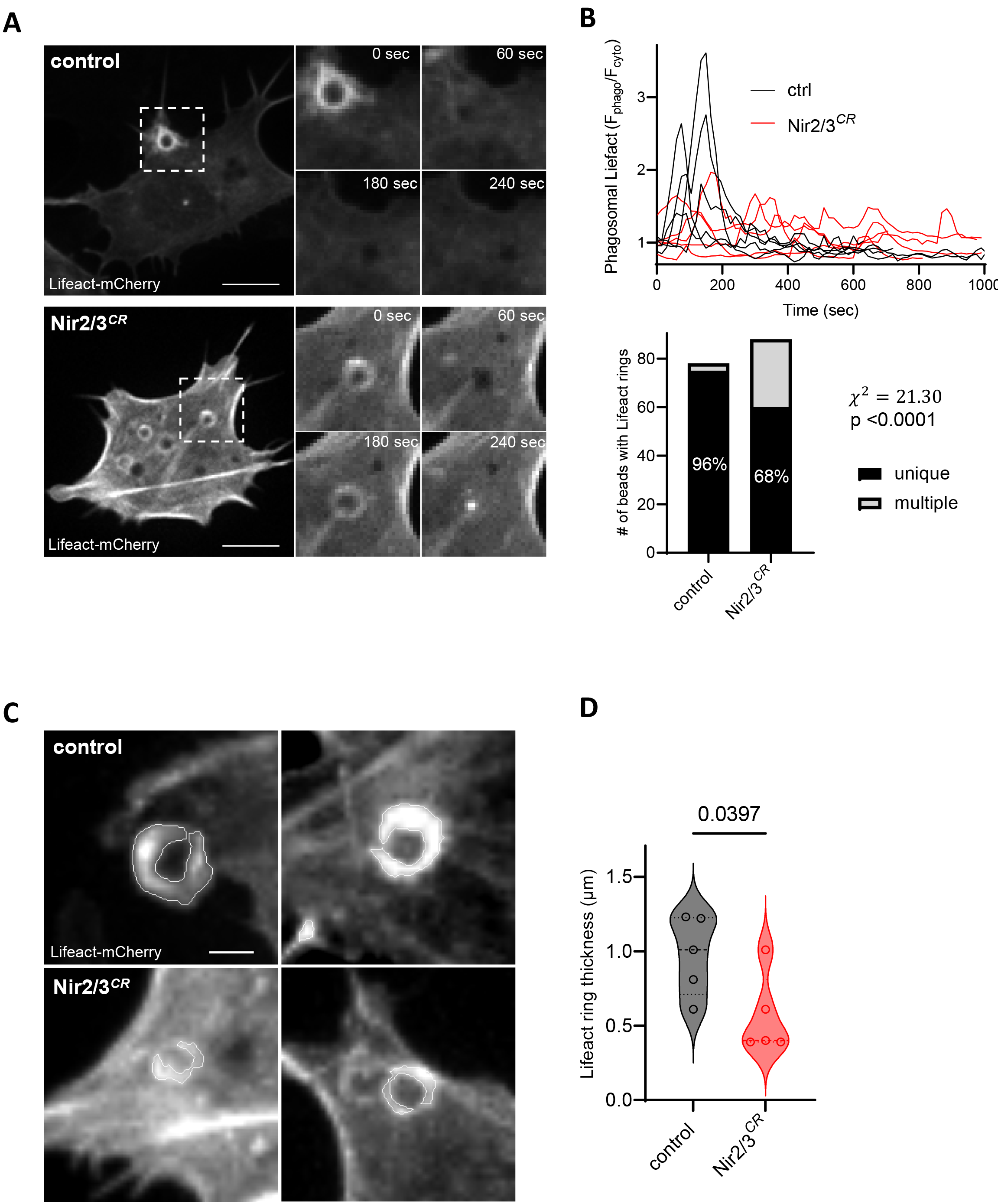
Nir2 and Nir3 depletion perturbs periphagosomal actin dynamics. A) Time-lapse images of LifeAct-mCherry dynamics during phagocytosis of IgG-opsonized particles by control (top) and Nir2/3-edited cells (bottom). Images from supplementary videos 1 and 2. Scale bars: 10 μm. B) Top: changes in phagosomal vs. cytosolic LifeAct-mCh fluorescence intensity during particle uptake by control and Nir2/3-edited cells (n=5 each). Bottom: proportion of unique vs. multiple occurrences of phagocytic LifeAct-mCh rings in control and Nir2/3-edited cells. C) Confocal sections at signal peak in control (top) and Nir2/3-edited cells (bottom). Lines indicate the spatial spread of the LifeAct-mCh accumulation. Scale bar: 2 μm. D) Thickness of periphagosomal LifeAct-mCh rings detected in control and Nir2/3-edited cells (n=5 each). Two-tailed unpaired t test.

## Discussion

Phosphoinositide metabolism regulates the efficiency of the phagocytic process initiated at particle capture and terminating at phagolysosome resolution (Botelho and Grinstein, 2011). The formation of phagocytic cups is associated with a transient local increase in PI(4,5)P2 that is quickly replaced by an equally transient elevation in PI(3,4,5)P3 at the site of contact, a biphasic change required for the remodeling of the actin cytoskeleton driving the extension of cups and the sealing of phagosomes. Both lipids promptly disappear from sealed phagosomes by the action of phosphatases to be replaced by PI(3,4)P2 and PI(3)P as phagosomes mature (Dewitt et al., 2006; Montaño-Rendón et al., 2022). ORP1L-mediated transfer of PI(4)P to the ER contributes to the resolution of phagosomes by promoting their tubulation (Levin-Konigsberg et al., 2019), but whether a lipid transfer protein is required for the supply of phosphoinositides at phagocytic cups is unclear as PI or PI(4)P can be delivered to cups by recycling endosomes (Levin-Konigsberg and Grinstein, 2020). Here, we show that the PI transfer proteins Nir2 and Nir3 participate in the lipid signaling events triggered by the engagement of phagocytic receptors by maintaining PI(4,5)P2 levels at sites of particle capture that enable the contractile activity of the actin cytoskeleton during phagosome sealing.

Using gene editing we show that combined depletion of Nir2 and Nir3 decreases basal PM PI(4,5)P2 levels and the efficiency of phagocytosis and that these two defects are corrected by re-expression of each of the two isoforms independently. This indicates that PM PI(4,5)P2 levels limit phagocytosis efficiency, consistent with earlier studies showing that the sequential phosphorylation of PI(4)P to PI(4,5)P2 and PI(3,4,5)P3 is required for phagosome formation (Montaño-Rendón et al., 2022). Interestingly, Nir2/3 editing blunted but did not completely eliminate dynamic elevations of PI(4,5)P2 during particle engulfment. This suggests that Nir proteins do not initiate but sustain the rapid amplification of the phosphoinositide signal generated by the activity of kinases. Therefore, once phagosomes are sealed, dynamic changes in phosphoinositides can occur within phagocytic membranes without Nir-mediated lipid exchange. In Nir2/3-edited cells, phagocytosis was halted at cup stage, indicating a reliance on Nir proteins for the sealing of phagosomes. Interestingly, Nir3 preferentially accumulated near phagocytic cups together with the ER-bound protein VAP-B, decorating ER cisternae vicinal to sealing phagosomes on the electron microscope. This suggests that Nir3 mediates the exchange of PA for PI mainly at phagocytic cups, a differential accumulation that might reflect the increased affinity of Nir3 for negatively charged lipids (Chang & Liou, 2015). Yet, each of the two lipid transfer proteins could restore phagocytic uptake when re-expressed, indicating that focal PI supply is not a strict requirement for cup closure, at least not in the context of copious global PM supply. We propose that both isoforms regulate phagocytic activity by maintaining PI signaling at sites of phagocytosis, with Nir2 acting predominantly at the PM and Nir3 at cups.

We previously showed that localized Ca^2+^ elevations can persist for tens of minutes around internalized phagosomes in primary mouse neutrophils and dendritic cells as well as MEF cells expressing FcgRIIa (Nunes et al., 2012). The long-lasting local Ca^2+^ elevations enhance the efficiency of phagocytosis and are fueled by Ca^2+^ release from InsP3R on juxtaphagosomal ER cisternae and by Ca^2+^ influx across STIM-gated Orai1 channels on phagosomes (Guido et al., 2015; Nunes et al., 2012). The persistent Ca^2+^ activity implies a constant local production of InsP3 around phagosomes to maintain the juxtaposed ER stores in a depleted condition enabling STIM1 to trap and gate phagosomal Orai1 channels. We therefore expected that Nir2/3 depletion would abort the local Ca^2+^ elevations by preventing the resupply of the PI consumed by signaling at ER-phagosome MCS. Contrary to our expectations, the Ca^2+^ signals persisted in cells depleted of Nir proteins. This indicates that Nir-mediated exchange of PA for PI is dispensable for the local generation of InsP3 and for the anchoring of STIM1 molecules to phagosomes once the particle has been engulfed. STIM1 binds to a range of PIPs via its polybasic domain and might remain bound to negatively charged lipids present on phagosomes until the resolution stage. Alternatively, STIM1 might be stabilized by junctate (Guido et al., 2015) or by binding to Orai1 channels on phagosomes. The constant generation of IP3 is more difficult to explain as the activity of PLC would stop without a constant supply of PI to regenerate its substrate PI(4,5)P2 in the membrane of phagocytic vacuoles. Other lipid transport proteins might substitute for Nir at the ER-phagosomes interface. Alternatively, the residual levels of PI(4,5)P2 might be sufficient for a small but sustained local production of IP3 enabling the opening of InsP3R clusters with a high sensitivity to IP3 immobilized at ER-phagosome MCS (Babu Thillaiappan et al., 2017). A pool of immobile IP3Rs licensed to respond to physiological stimuli localizes near STIM-ORAI interaction sites at ER-PM junctions (Taylor and Machaca, 2019). Whether this mechanism sustains prophagocytic Ca^2+^ elevations at ER-phagosome MCS remains to be established.

A striking phenotype of Nir2/3-edited cells was the abnormal pattern of actin condensation occurring around forming phagosomes. The thickness of the dense F-actin rings surrounding sites of particle capture was significantly below the levels observed in control cells, and multiple occurrences of ring formation were observed in apparent unsuccessful attempts of phagosome closure. This phenomenon was not reported previously and likely reflects the deficiency in local non-vesicular lipid transport leading to reduced PI(4,5)P2 levels at cups. A substantial amount of evidence indicates that PI(4,5)P2 accumulation at cups recruits and activates actin-modulating proteins to promote F-actin accumulation and contractile activity at cups (Raucher et al., 2000; Yeung and Grinstein, 2007). Reduced F-actin amounts at phagosomal cups were reported in macrophages expressing a kinase-dead mutant of phosphatidylinositol-4-phosphate 5-kinase (PIPK), the enzyme that generates PI(4,5)P2 from PI(4)P at cups (Coppolino et al., 2002). PIPK-edited macrophages exhibited normal particle binding and receptor clustering but reduced accumulation of PI(4,5)P2 at cups and phagocytosis was blocked at cup stage, as we report here for Nir2/3-deficiency. In contrast, manipulations that enhance PI(4,5)P(2) levels at cups cause persistent actin accumulation (Scott et al., 2005). The loss of Nir proteins therefore likely halts phagocytosis at cup stage because the decreased local PI(4,5)P2 levels prevent the accumulation of cortical actin at cups, decreasing the contractile activity required for particle internalization.

In conclusion, we show that phosphatidylinositol transfer mediated by Nir2/3 proteins is required for the engulfment of phagocytic targets. Genetic disruption in the genes encoding for these proteins reduces basal PI(4,5)P2 levels in phagosomal membranes and prevents the formation of contractile actin rings around captured particles, aborting phagocytosis at cup stage without impacting PI(4,5)P2 and Ca^2+^ elevations during phagosome maturation. Nir2 preferentially mediates lipid exchange at the plasma membrane and Nir3 at phagocytic cups but either isoform can independently complement the phagocytic defect caused by the combined depletion of both proteins. We conclude that PI(4,5)P2 levels at cup stage are limiting for the recruitment of the dense cortical actin cytoskeleton driving phagosome sealing.

## Material and methods

### Antibodies and reagents

Mouse embryonic fibroblasts (MEF) were kindly provided by Luca Scorrano (Padova, Italy) and regularly tested for mycoplasma contamination. Myc-tagged human FcγRIIA was gifted by Sergio Grinstein (Toronto, Canada), EGFP-Nir2 and EGFP-Nir3 by Tamas Balla (NIH, USA), LifeAct-mCherry by Florence Niedengang (Institut Cochin, Paris). PH-PLCδ1-GFP (#51407) and mCherry-VAP-B (#108126) were purchased from Addgene, pTagRFP-C (#FP141) from Evrogen, fura-2-AM, BAPTA-AM, Lipofectamine 2000 from Life Technologies. TagRFP-Nir2 and TagRFP-Nir3 were subcloned from EGFP-Nir2 and EGFP-Nir3 into pTagRFP-C, respectively. AF633 coupled goat anti-human IgG (#A-21091) was purchased from Thermo Fisher, rat anti-mouse CD16/32 (Mouse BD Fc Block, #553142) from BD Biosciences, anti-human CD32-APC antibody from Miltenyi Biotec. Ca^2+^ recordings and live imaging experiments were conducted in physiological buffer containing 140 mM NaCl, 5 mM KCl, 1 nM MgCl_2_, 2 mM CaCl_2_, 20 mM Hepes, 10 mM glucose, pH 7.4 with NaOH.

### Cell lines, cell culture, transfection and transduction

MEF cells were cultured in Dulbecco’s modified Eagle’s medium (DMEM; high glucose, catalog number 31966, Life Technologies) containing 10% fetal calf serum (FCS) and 0.5% penicillin-streptomycin (pen-strep, catalog number 15140, Life Technologies) at 37°C under 5% CO2 and were passaged twice a week. CRISPR MEF cells were generated using Double Nickase Plasmid (Santa Cruz Biotechnology) against mouse *Nir2* and *Nir3* genes. Cells were transfected using Lipofectamine 2000 at 40-50% confluence in high glucose (4,5 g/l) DMEM without serum, selected with puromycin, and sorted on GFP fluorescence. Sorted single cell clones were screened for Nir2 and Nir3 depletion by genomic PCR. Absence of the product amplified using the primer binding to the expected cut site suggested a modification of the sequence which was validated by sequencing of the region including the cut site. qPCR was also performed to validate the decrease in mRNA levels. TagRFP-C1, TagRFP-Nir2 and TagRFP-Nir3 were subcloned into pLenti-CMVie-IRES-BlastR (#119863, Addgene) and constructs were co-transfected with pMD2G/psPAX2 into HEK-293T cells to produce viral particles as described previously (Carreras-Sureda et al., 2021). After transduction in the presence of polybrene at 10 μg/ml, stable cells were obtained by antibiotic selection (blasticidin, 5 μg/ml) based on TagRFP fluorescence by FACS sorting.

### Phagocytic target preparation

Carboxyl polystyrene microsphere (3.0 μm, Spherotech, CP-30-10) were opsonized by covalent coupling with hIgG. Following three washes in sterile PBS at 10000g at 4°C for 3 min, polystyrene beads were activated by 50 mM 1-Ethyl-3-(3-dimethylaminopropyl) carbodiimide hydrochloride (Carl Roth/2156.1) in PBS for 15 min at room temperature by shaking. Next, beads were washed 3 times in 0.1 M Na2B4O7 buffer (pH 8.0) at 4°C and 6 mg of hIgG were added to beads and incubated at 4°C overnight on a shaker. To prepare fluorescently labeled hIgG polystyrene beads, Alpha Fluor 488 amine (20 μg/mL, AAT Bioquest/Cat No. 1705) was added at room temperature for 30 min with agitation before overnight incubation with hIgG. Beads were washed twice with 250 mM glycine/PBS and then with PBS alone, resuspended in sterile PBS and 0.5% pen/strep and counted by a cell counter.

### Phagocytic activity assessment

AF-488 hIgG beads were added at target:cell ratio of 10:1 and mildly centrifuged onto cells. Following incubation at 37°C under 5% CO_2_ for indicated times in serum-containing media, cells were washed and blocked with cold FACS buffer (1% BSA, 5 mM EDTA in PBS) for 15 min at 4°C. Next, cells were incubated with APC-CD32 antibody to determine the Fc positive population. Flow cytometry measurements were performed on a BDLSR Fortessa (Becton Dickinson) and analyzed using FlowJo software.

### Calcium imaging

Periphagosomal Ca^+2^ microdomain measurement imaging was conducted as indicated (Guido et al., 2015; Nunes et al., 2012). Fluo8-AM (4 μM, AAT Bioquest) was loaded in physiological buffer with sulfinpyrazone, for 30 min first at 37°C, then 20 min at RT and with BAPTA-AM (2,5 μM) for 10 more minutes. Fluorescently labelled IgG-opsonized targets were mildly centrifuged onto cells and incubated for 30 min. Images were acquired on a Nipkow Okagawa Nikon spinning disk confocal microscopy with a temperature controller, motorized stage and Plan Apo 40x/1.3 Oil DICIII objective, using Visiview software (Visitron Systems). At least five snapshots, averaged over 6s, per coverslip were quantified by custom ImageJ macros as described previously (Guido et al., 2015; Nunes et al., 2012). Briefly, local Ca^+2^ elevations were defined as microdomains if the area is ≥500 nm^2^ within a distance of ∼750 nm from the phagosome membrane with a fluorescence value at least 2 x s.d higher than the average cytoplasmic Fluo8 intensity.

### Live Imaging

All the live imaging experiments were performed on a Nikon spinning disk confocal microscopy with a temperature controller, motorized stage and Plan Apo 40x/1.3 and Plan Apo 63x/1.4 Oil DICIII objective using Visiview software (Visitron Systems). Cells were seeded on 25 mm coverslips the day before transfection and mounted on AttoFluor chambers for microscopy with 2 mM Ca^2+^ medium before the experiment. For phagosomal lipid assessment, IgG-opsonized targets were added 5-10 min before starting recording. At least 5 stage positions were selected with cells already started phagocytosing. Z-stack of 9-12 slices spaced at 0.5μm were imaged every 30s for each stage position during 20-30 min. Sum projections of background-subtracted images and individual phagosome tracking were done manually with ImageJ. Phagosomal PI(4,5)P2 and actin enrichment was determined by the ratio of the average signal from the phagosome membrane to local average cytosolic signal for the corresponding time point. For PM PI(4,5)P2 analysis, z-stack of 6-8 slices spaced at 0.5μm were acquired on 7-10 stage positions per coverslip. Background-subtracted, average projection images of three central slices were created by ImageJ. Ratio of PM fluorescence over cytosol fluorescence was used to determine the enrichment at the PM.

### Immunolabelling

Cells were fixed in 4% paraformaldehyde (PFA) in PBS for 20 min at room temperature, permeabilized in PBS-BSA 0.5 %+ NP-40 0.1% for 10 min and blocked in PBS-BSA 0.5 %+ FBS 5% for 1 hour at room temperature. For labeling of external beads, after fixation cells were blocked in Fc block in PBS-BSA 1% (1:200) and incubated with anti-human IgG antibody in PBS-BSA 0.5% (1:500) for 1 hour at room temperature. Then, cells were incubated with primary antibodies (Myc-Tag, 9B11, Cell Signaling) overnight in a humid chamber at 4°C. Next day, cells were incubated with secondary antibody (1:5000) with Hoesch (1:10000) for 1 hour at room temperature. Coverslips were mounted in Fluoromount-GT Slide Mounting Medium (Electron Microscopy Sciences). Beads at cup stage were quantified as the number of beads labeled with human IgG and in contact with cells expressing Fc receptor divided by the total number of beads (internalized + cup stage). Phagocytic index was determined as the number of ingested beads divided by total number of Fc receptor expressing cells.

### Correlation Light Electron Microscopy (CLEM)

Focused-ion beam scanning electron microscopy (FIB-SEM) was performed on a Helios NanoLabG3 microscope (FEI. Netherlands), as described (Nunes-Hasler et al., 2017). Briefly, transfected cells were seeded on 35-mm Ibidi polymer dishes with gridded bottom (catalog number 81166). Following incubation with the target and fixation with PFA 4% for 20 min at room temperature, brightfield and high-resolution confocal images were captured. Next, samples were fixed for EM in 2.5% glutaraldehyde/2% PFA/2 mM CaCl_2_/0.15 M sodium cacodylate buffer (Ca-Caco) at pH 7.4 for 3 hours on ice and washed 5x in ice cold Ca-Caco. Following dehydration, samples were embedded on Epon and prepared for FIB-SEM imaging as previously described. Samples were sputter-coated with gold for 30 seconds by a Q150T ES coater (Quorum Technologies, UK). Cellular footprints obtained from FIB-SEM and fluorescence and brightfield images were compared to locate the cell of interest. Images were acquired at the highest resolution setting (5x5x10 nm) using the Autoslice and View software (FEI). Drift correction and alignment of FIB-SEM images were done using Amira Software and overlay with the fluorescence image using ImageJ.

### Image analysis and statistics

All images were analyzed with ImageJ software. All statistical analyses were performed with GraphPad Prism 9. Unless otherwise indicated all statistical tests conducted are two-tailed.

## Acknowledgements

We thank Cyril Castelbou for the technical assistance and the bioimaging and electron microscopy core facilities of the Faculty of medicine of the University of Geneva. This work was funded by the Swiss National Foundation [grant number 310030_189042 (to N.D.) and 310030_189094 (to P.N.H.)], the Sir Jules Thorn Charitable Trust Foundation (to A.C.S) and the Novartis foundation (to A.C.S) P.N.H is recipient of a career award from the Prof. Dr. Max Cloëtta Foundation.

## Data availability

The data that support the findings of this study are presented in the main and supplementary figures, with number of experiments and statistical tests applied. Primary data are available from the corresponding author upon reasonable request.

## Author contributions

MK, Acquisition of data, Analysis and interpretation of data, Drafting or revising the article; ACS, PN, & ND, Conception and design, Analysis and interpretation of data, Drafting or revising the article

## Figure legends

**Figure S1.**
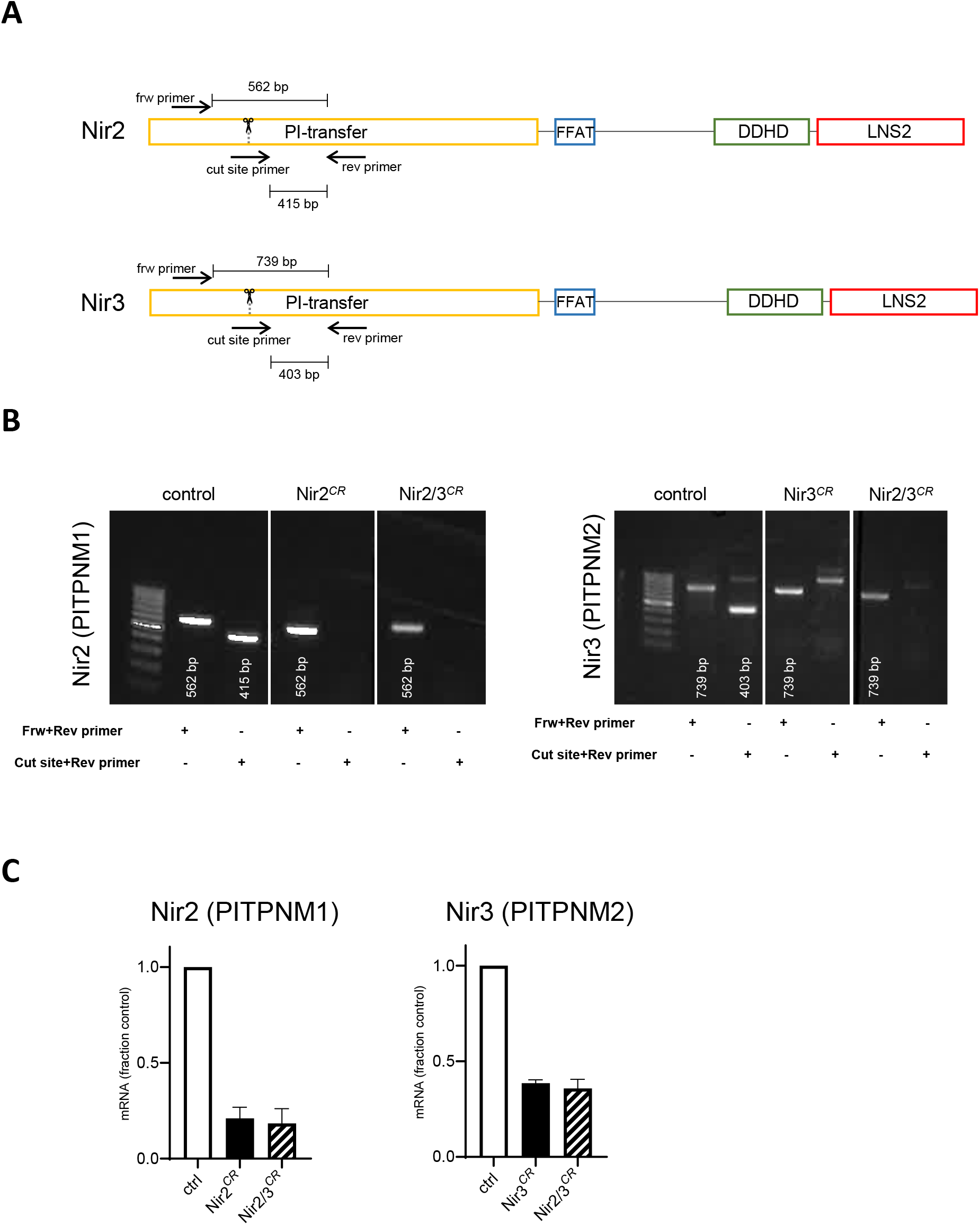
Related to Fig. 2. Genetic and functional validation of Nir2/3 depletion. A) Sites in the PITPNM genes targeted by the guide RNAs and PCR primers. B) PCR products detected by primers targeting regions outside or within the cut site in the indicated clones. C) Nir2 (top) and Nir3 (bottom) mRNA expression in the indicated clones relative to CRISPR-control cells (n=2).

**Figure S2.**
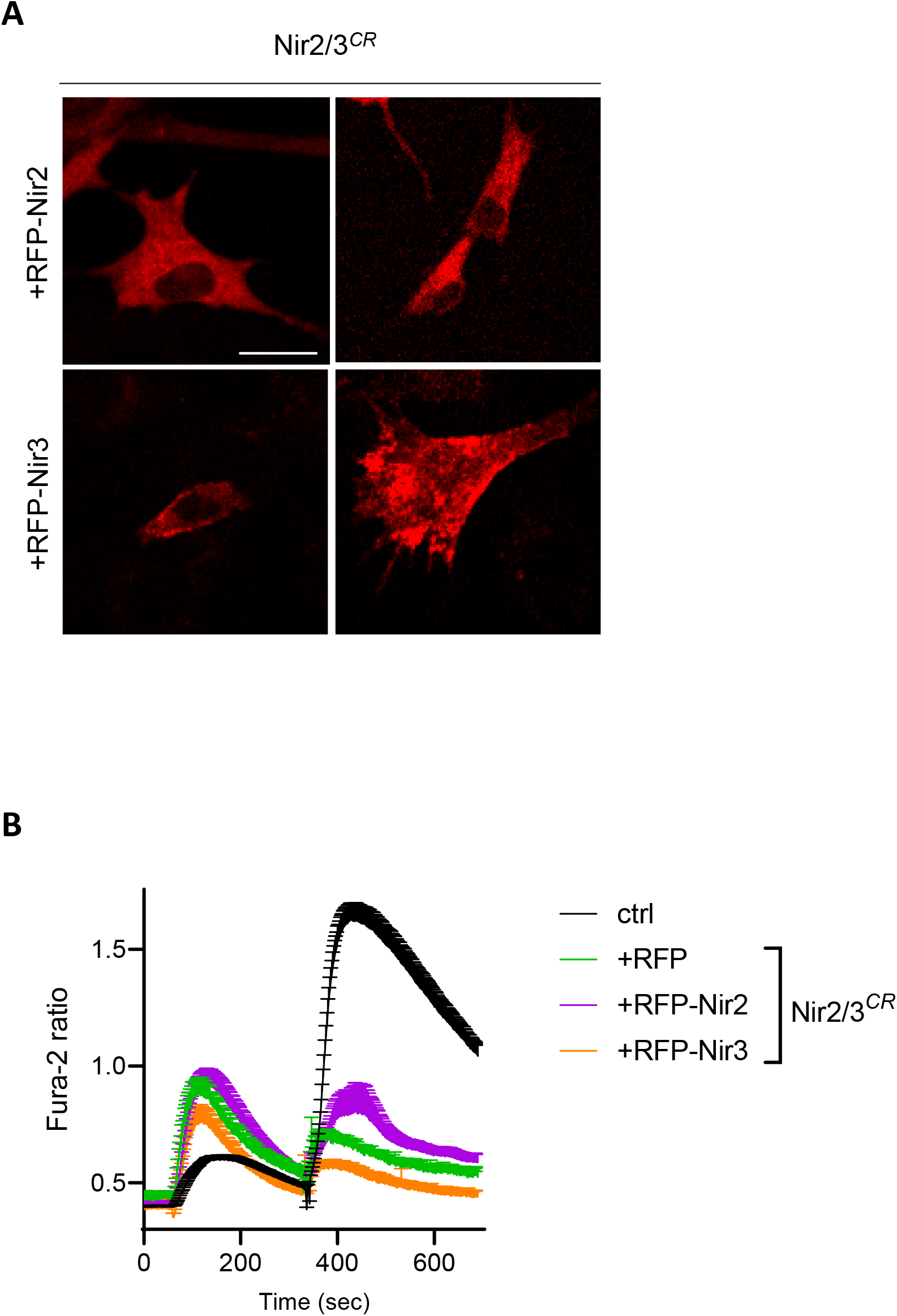
Related to Fig. 2. RFP-Nir2 and RFP-Nir3 re-expression does not rescue SOCE. A) Fluorescence images of Nir2/3-edited cells expressing the indicated RFP-tagged constructs. Cells were imaged on a wide-field microscope at λex:560 nm, λem:633 nm. B) Averaged Ca^2+^ responses evoked by the addition of thapsigargin and 1 mM Ca^2+^ to Nir2/3-edited cells expressing the indicated constructs. Recordings totaling 11-35 cells.

**Figure S3.**
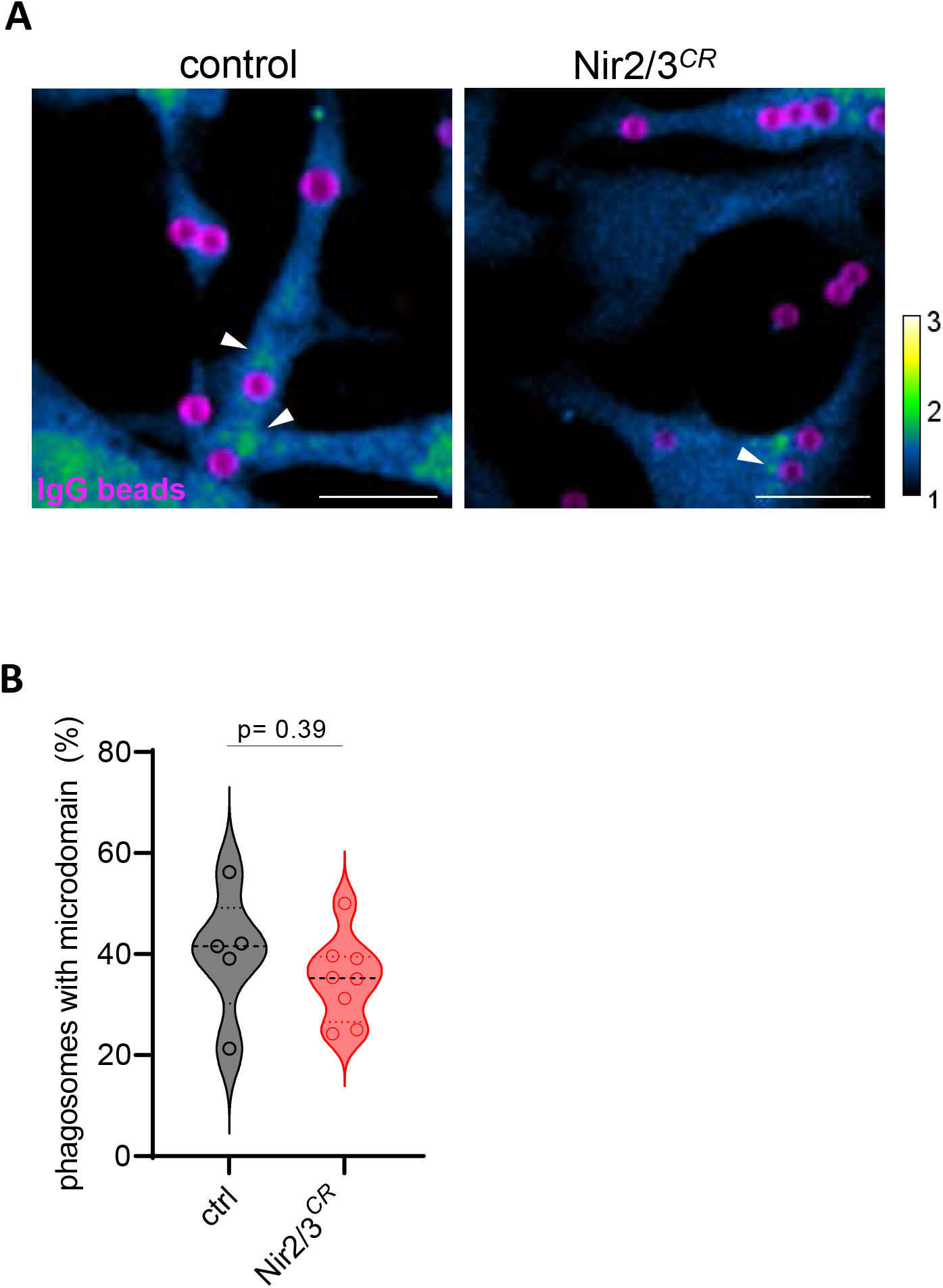
Related to Fig. 3. Effect of Nir2/3 depletion on periphagosomal calcium signals. A) Fluo-8 ratio images (F/F_0_) of control and Nir2/3-edited cells loaded with 4 μM Fluo8-AM and 2.5 μM BAPTA-AM in medium containing 2mM Ca^2+^ after 30 min of phagocytosis. Scale bars 10 μm. B) Proportion of phagosomes associated with periphagosomal Ca^2+^ hotspots in control and Nir2/3-edited cells. Two-tailed unpaired t test of n=6 and 8 recordings with 96/265 and 91/253 cells/phagosomes each for control and Nir2/3-edited cells, respectively.

**Figure S4.**
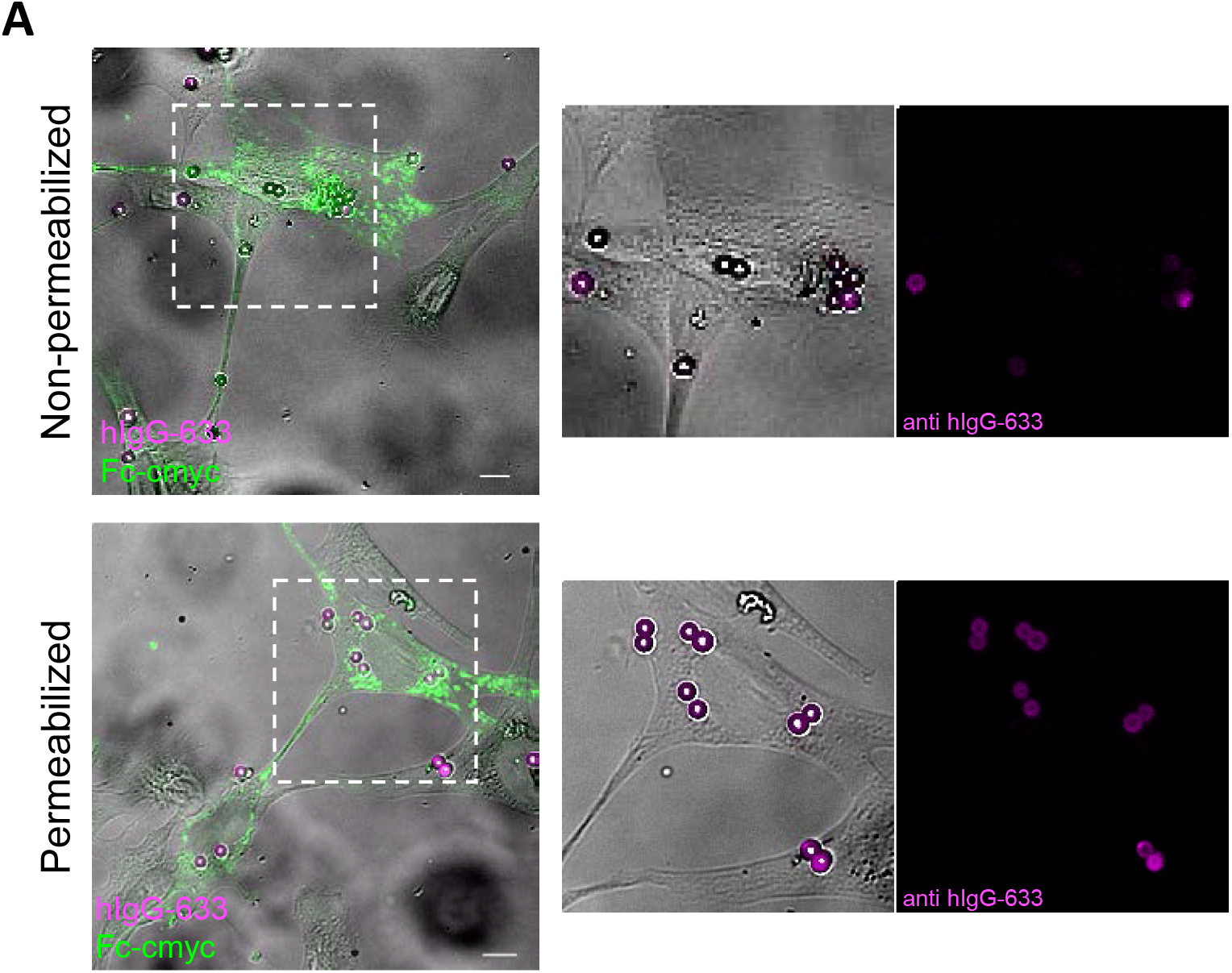
Related to Fig. 4 -Nir2 and Nir3 depletion stalls phagocytosis at cup stage. Bright-field and fluorescence images of cells expressing myc-FcRIIA labeled with anti-cMyc (green) and anti-hIgG (magenta) permeabilized (bottom) or not (top) with NP40 0.01% 30 min after particle exposure. Scale bars: 10 μm.

## References

Amarilio, R., Ramachandran, S., Sabanay, H. and Lev, S. (2005). Differential Regulation of Endoplasmic Reticulum Structure through VAP-Nir Protein Interaction. J. Biol. Chem. 280, 5934–44.

Babu Thillaiappan, N., Chavda, A. P., Tovey, S. C., Prole, D. L., Taylor, C. W. and And, R. (2017). Ca2+ signals initiate at immobile IP 3 receptors adjacent to ER-plasma membrane junctions. Nat. Commun.

Balla, T. (2013). Phosphoinositides: Tiny Lipids With Giant Impact on Cell Regulation. Physiol Rev 93, 1019–1137.

Balla, T. (2018). Ca2+ and lipid signals hold hands at endoplasmic reticulum–plasma membrane contact sites. J. Physiol. 596, 2709–2716.

Balla, T. and Várnai, P. (2002). Visualizing cellular phosphoinositide pools with GFP-fused protein-modules. Sci. STKE 2002, 1–17.

Bohdanowicz, M. and Grinstein, S. (2013). Role of phospholipids in endocytosis, phagocytosis, and macropinocytosis. Physiol. Rev. 93, 69–106.

Botelho, R. J. and Grinstein, S. (2011). Phagocytosis. Curr. Biol. 21,.

Botelho, R. J., Teruel, M., Dierckman, R., Anderson, R., Wells, A., York, J. D., Meyer, T. and Grinstein, S. (2000). Localized biphasic changes in phosphatidylinositol-4,5-bisphosphate at sites of phagocytosis. J. Cell Biol. 151, 1353–1367.

Carreras-Sureda, A., Abrami, L., Kim J.H., Wang, W.A., Henry, C., Frieden, M. Didier, M. van der Goot, F.G. and Demaurex, N. (2021). S-acylation by ZDHHC20 targets ORAI1 channels to lipid rafts for efficient Ca2+ signaling by Jurkat T cell receptors at the immune synapse. Elife 10:e72051.

Chang, C. L. and Liou, X. J. (2015). Phosphatidylinositol 4,5-Bisphosphate Homeostasis Regulated by Nir2 and Nir3 Proteins at Endoplasmic Reticulum-Plasma Membrane Junctions. J. Biol. Chem. 290, 14289–14301.

Chang, C. L. and Liou, J. (2016). Homeostatic regulation of the PI(4,5)P2-Ca2+ signaling system at ER-PM junctions. Biochim. Biophys. Acta - Mol. Cell Biol. Lipids 1861, 862–873.

Cockcroft, S. and Garner, K. (2011). Function of the phosphatidylinositol transfer protein gene family: is phosphatidylinositol transfer the mechanism of action? Crit Rev Biochem Mol Biol. 46, 89–117

Coppolino, M. G., Dierckman, R., Loijens, J., Collins, R. F., Pouladi, M., Jongstra-Bilen, J., Schreiber, A. D., Trimble, W. S., Anderson, R. and Grinstein, S. (2002). Inhibition of phosphatidylinositol-4-phosphate 5-kinase Ia impairs localized actin remodeling and suppresses phagocytosis. J. Biol. Chem. 277, 43849–57.

Dewitt, S., Tian, W. and Hallett, M. B. (2006). Localised Ptdlns(3,4,5)P3 or Ptdlns(3,4)P2 at the phagocytic cup is required for both phagosome closure and Ca2+ signalling in HL60 neutrophils. J. Cell Sci. 119, 443–451.

Flannagan, R. S., Jaumouillé, V. and Grinstein, S. (2012). The Cell Biology of Phagocytosis. Annu. Rev. Pathol. Mech. Dis 7, 61–98.

Grinstein, S. (2016). Multiphasic dynamics of phosphatidylinositol 4-phosphate during phagocytosis. Mol. Biol. Cell 28, 128–140.

Guido, D., Demaurex, N. and Nunes, P. (2015). Junctate boosts phagocytosis by recruiting endoplasmic reticulum Ca 2+ stores near phagosomes. J. Cell Sci. 128, 4074–82.

Kang, F., Zhou, M., Huang, X., Fan, J., Wei, L., Boulanger, J., Liu, Z., Salamero, J., Liu, Y. and Chen, L. (2019). E-syt1 Re-arranges STIM1 Clusters to Stabilize Ring-shaped ER-PM Contact Sites and Accelerate Ca 2+ Store Replenishment. Sci. Rep. 9:3975.

Keinan, O., Kedan, A., Gavert, N., Selitrennik, M., Kim, S., Karn, T., Becker, S. and Lev, S. (2014). The lipid-transfer protein Nir2 enhances epithelial-mesenchymal transition and facilitates breast cancer metastasis. J. Cell Sci. 127, 4740–4749.

Kim, Y. J., Guzman-Hernandez, M. L., Wisniewski, E., Echeverria, N. and Balla, T. (2016). Phosphatidylinositol and phosphatidic acid transport between the ER and plasma membrane during PLC activation requires the Nir2 protein. Biochem Soc Trans. 44, 197–201.

Levin-Konigsberg, R. and Grinstein, S. (2020). Phagosome-endoplasmic reticulum contacts: Kissing and not running. Traffic 21, 172–180.

Levin-Konigsberg, R., Montaño-Rendón, F., Keren-Kaplan, T., Li, R., Ego, B., Mylvaganam, S., Diciccio, J. E., Trimble, W. S., Bassik, M. C., Bonifacino, J. S., et al. (2019). Phagolysosome resolution requires contacts with the endoplasmic reticulum and phosphatidylinositol-4-phosphate signalling. Nat. Cell Biol. 21, 1234–1247.

Marshall, J. G., Booth, J. W., Stambolic, V., Mak, T., Balla, T., Schreiber, A. D., Meyer, T. and Grinstein, S. (2001). Restricted Accumulation of Phosphatidylinositol 3-Kinase Products in a Plasmalemmal Subdomain during Fc Receptor-mediated Phagocytosis. J. Cell Biol. 153, 1369–80.

Montaño-Rendón, F., Walpole, G. F., Krause, M., Hammond, G. R., Grinstein, S. and Fairn, G. D. (2022). PtdIns(3,4)P2, Lamellipodin, and VASP coordinate actin dynamics during phagocytosis in macrophages. J. Cell. Biol. 221:e202207042.

Nunes-Hasler, P., Maschalidi, S., Lippens, C., Castelbou, C., Bouvet, S., Guido, D., Bermont, F., Bassoy, E. Y., Page, N., Merkler, D., et al. (2017). STIM1 promotes migration, phagosomal maturation and antigen cross-presentation in dendritic cells. Nat. Commun.

Nunes-Hasler, P., Kaba, M. and Demaurex, N. (2020). Molecular Mechanisms of Calcium Signaling During Phagocytosis. Adv. Exp. Med. Biol. 1246, 103–128.

Nunes, P., Cornut, D., Bochet, V., Hasler, U., Oh-Hora, M., Waldburger, J. M. and Demaurex, N. (2012). STIM1 juxtaposes ER to phagosomes, generating Ca2+ hotspots that boost phagocytosis. Curr. Biol. 22, 1990–1997.

Raucher, D., Stauffer, T., Chen, W., Shen, K., Guo, S., York, J. D., Sheetz, M. P. and Meyer, T. (2000). Phosphatidylinositol 4,5-bisphosphate functions as a second messenger that regulates cytoskeleton-plasma membrane adhesion. Cell 100, 221–228.

Scott, C. C., Dobson, W., Botelho, R. J., Coady-Osberg, N., Chavrier, P., Knecht, D. A., Heath, C., Stahl, P. and Grinstein, S. (2005). Phosphatidylinositol-4,5-bis phosphate hydrolysis directs actin remodeling during phagocytosis. J. Cell Biol. 169, 139–149.

Selitrennik, M. and Lev, S. (2016). The role of phosphatidylinositol-transfer proteins at membrane contact sites. Biochem. Soc. Trans. 44, 419–424.

Taylor, C. W. and Machaca, K. (2019). IP 3 receptors and store-operated Ca 2+ entry: a license to fill. Curr. Opin. Cell Biol. 57, 1–7.

Uribe-Querol, E. and Rosales, C. (2020). Phagocytosis: Our Current Understanding of a Universal Biological Process. Front. Immunol. 11, 1066. doi: 10.3389/fimmu.2020.01066.

Várnai, P., Rother, K. I. and Balla, T. (1999). Phosphatidylinositol 3-kinase-dependent membrane association of the Bruton’s tyrosine kinase pleckstrin homology domain visualized in single living cells. J. Biol. Chem. 274, 10983–10989.

Westman, J., Grinstein, S. and Maxson, M. E. (2019). Revisiting the role of calcium in phagosome formation and maturation. J. Leukoc. Biol. 1–15.

Yeung, T. and Grinstein, S. (2007). Lipid signaling and the modulation of surface charge during phagocytosis. Immunol. Rev. 219, 17–36.

Yeung, T., Ozdamar, B., Paroutis, P. and Grinstein, S. (2006). Lipid metabolism and dynamics during phagocytosis. Curr. Biol. Cell Biol. 18, 429–437.

